# Single Nucleus Transcriptomics Reveals Pervasive Glial Activation in Opioid Overdose Cases

**DOI:** 10.1101/2023.03.07.531400

**Authors:** Julong Wei, Tova Y. Lambert, Aditi Valada, Nikhil Patel, Kellie Walker, Jayna Lenders, Carl J. Schmidt, Marina Iskhakova, Adnan Alazizi, Henriette Mair-Meijers, Deborah C. Mash, Francesca Luca, Roger Pique-Regi, Michael J Bannon, Schahram Akbarian

## Abstract

Dynamic interactions of neurons and glia in the ventral midbrain (VM) mediate reward and addiction behavior. We studied gene expression in 212,713 VM single nuclei from 95 human opioid overdose cases and drug-free controls. Chronic exposure to opioids left numerical proportions of VM glial and neuronal subtypes unaltered, while broadly affecting glial transcriptomes, involving 9.5 - 6.2% of expressed genes within microglia, oligodendrocytes, and astrocytes, with prominent activation of the immune response including interferon, NFkB signaling, and cell motility pathways, sharply contrasting with down-regulated expression of synaptic signaling and plasticity genes in VM non-dopaminergic neurons. VM transcriptomic reprogramming in the context of opioid exposure and overdose included 325 genes with genetic variation linked to substance use traits in the broader population, thereby pointing to heritable risk architectures in the genomic organization of the brain’s reward circuitry.

## INTRODUCTION

Ventral midbrain (VM), including the ventral tegmental area (VTA) and substantia nigra (SN)^1,2^, is important for mediating habitual behaviors and salience of cues associated with drug use, as well as withdrawal-related anhedonia and dysphoria^3,4^. It has become increasingly clear in recent years that, in addition to the well-established roles of DA and non-DA (e.g., GABAergic) neurons, the VM’s glial and other non-neuronal populations may play an important role for drug responsiveness and substance use. To mention just three representative examples, excessive activation of VM microglia is thought to disrupt chloride homeostasis in GABA neurons, which in turn, negatively affects opioid- and stimulant-induced dopamine release and associated reward behaviors^5^. Likewise, VM astrocytes play an essential role in drug-induced synaptic plasticity in DA neurons^6^, a reflection of astrocytic regulation of neuronal glutamine supply and glutamatergic neurotransmission^7^. Finally, oligodendrogenesis in VM is essential for morphine-mediated reward behavior, and proliferation and differentiation of VM oligodendrocytes is regulated by the firing activity of their surrounding dopaminergic neurons, in addition to direct effects exerts by activated opioid receptors expressed on the oligodendrocytes^8^.

However, despite these intriguing mechanistic studies in animal models, the functional and clinical significance of VM glial populations in subjects diagnosed with substance use disorder remains unexplored. To this end, cell-specific transcriptomic profiling of VM dissected from human post-mortem brain could deliver critical insights. This task is particularly urgent for opioid use disorder (OUD), considering that opioid overdose (OD) is now the leading cause of accidental deaths in the United States, with ∼70,000 deaths annually reflecting a > 8-fold increase over the course of just two decades^9^. However, to date, with the exception of a single study profiling RNA from VM bulk tissue in a limited cohort of cases and controls^10^, no knowledge exists about genome-scale dysregulation associated with chronic opioid exposure and overdose. Further, RNA-seq profiling of VM bulk tissue is insufficient to disentangle cell type-specific contributions in neuropsychiatric disease^11^.

Of note, recent pilot studies exploring adult postmortem human VM single cell genomics (by 10x chromium single nuclei transcriptomic profiling; Smajic and colleagues^12^, and others^13^, reported very high recovery rates of glial and other non-neuronal nuclei in these ‘dopaminergic’ brain structures, with >95-96% of the total population of nuclei recovered from the SN contributed by prototypical glia, including oligodendrocytes and their precursors, astrocytes, and microglia. In contrast, DA and GABA neurons taken together contributed only a very minor (< 4%) share of nuclei in this type of single nuclei RNA-seq assay^12,13^. This approach thus lends itself to an in-depth characterization of opioid-related changes in gene expression in these relatively under-studied glial cell types in human VM.

Here, we present our findings from a transcriptomic study at single nuclei resolution in the VM, built from two independent case-control cohorts (totaling 95 subjects) from different geographical areas in the U.S. We report, for the first time, highly reproducible alterations affecting hundreds of microglia-, astrocyte- and oligodendroglia-associated transcripts. Remarkably, these widespread, cell type-specific disruptions of the glial VM transcriptome in opioid overdose cases occurred in the context of completely conserved cellular composition, with stoichiometric proportions for all neuronal and glial subtypes indistinguishable between cases and controls. Our findings point to profound alterations of gene expression in victims of opioid overdose, indicative of neuroinflammation and activation of cytokine signaling in the VM, primarily affecting microglia and astrocytes, with additional alterations of oligodendrocyte-specific transcriptomes. More broadly, the dataset presented here will provide a much needed human neurogenomics resource at single cell resolution for the wider field of drug abuse research.

## RESULTS

### Chronic opioid exposure does not alter the cellular composition of the ventral midbrain

We generated VM single nuclei RNA-seq libraries for 95 brain donors (84M/11F), including 45 subjects with documented histories of opioid abuse and overdose, and 50 demographically-matched, opioid-free control subjects, collected from two geographically distinct regions within the U.S. (greater Detroit area, Michigan and Miami, Florida) (***Figure 1A, Table S1***). Each VM sample included both substantia nigra and the adjacent ventral tegmental area (SN/VTA) (see Methods). Nuclei were processed in pools of 3-4 brains using the 10X Chromium system followed by Illumina sequencing, read alignment and processing by 10X Cellranger. Each single nucleus was matched to a donor using Demuxlet, confirming a 100% match by donor by pool against the background of all 95 donors (***Figure S1A***). After removal of doublets and quality-control filtering (see Methods), we obtained a total of 212,713 transcriptionally profiled single nuclei, each unique to a singular donor (median, 2,008 nuclei/donor). We collected 2,696∼21,363 (median, 8,274) reads/nucleus (***Table S2***) and measured the expression of 1,383∼5,079 (median, 3,070) genes/nucleus. Total numbers of single nuclei/specimen, genes called/single nucleus/specimen and read depth/nucleus/specimen showed no significant differences between cases and controls (***Figure S1B-D***).

**Figure 1:**
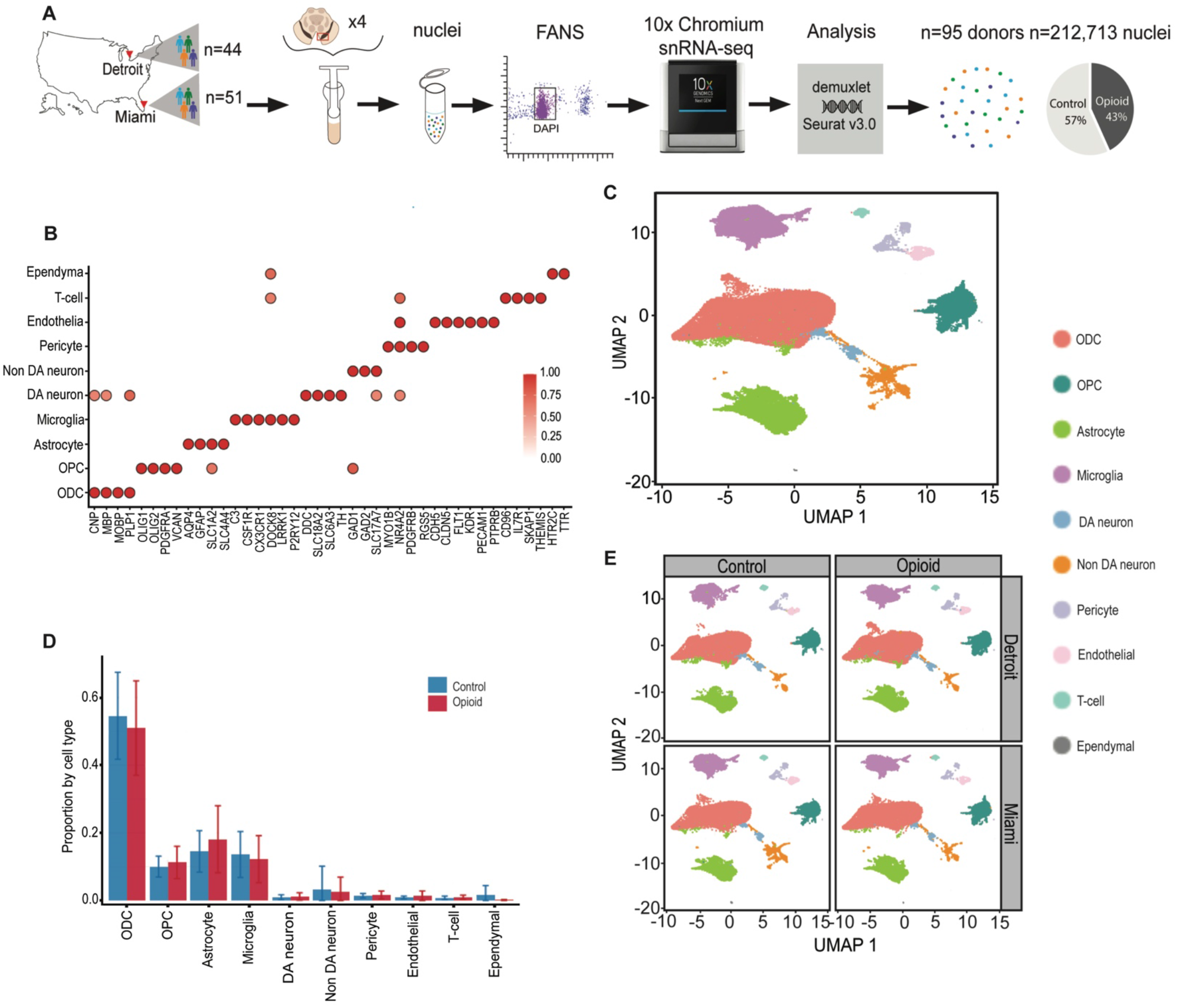
Consistent proportions of VM cell types across specimen collection sites and cohorts. (**A**) Experimental design and workflow for N=95 VM samples, including collection at two different geographical areas, pooling of 3-4 VM/substantia nigra specimens, nuclei purification by FACS and 10x chromium snRNA-seq pipeline and genetic fingerprinting of nuclei by donor, yielding a total of 212,713 nuclei. (**B**) Marker gene expression for each of the 10 glial and neuronal subpopulations as indicated. DA, dopaminergic neuron. non-DA, non-dopaminergic neurons; ODC oligodendrocyte, OPC, oligodendrocyte precursor cell. Color represents the ratio of average gene expression across cells in the cell type relative to maximum in the most highly enriched in some cell-type.(**C**) Uniform Manifold Approximation and Projection (UMAP) plot showing identified 10 major cell types by cluster, as indicated, for total collection of n=212,713 nuclei. (**D**) Proportional representation of cell types across individual donors (mean and S.D. of each of the 10 major cell types by diagnosis (red opioid-related death, blue control) as indicated. (**E**) VM cell type composition by UMAP plot, shown for cases and controls separately for each of the two collection sites.

Resolving the entire collection of 212,713 nuclei by cluster analysis in Seurat v.4.0 with 2,000 highly variable genes and 50 harmony-adjusted principal components produced in the Uniform Manifold Approximation and Projection (*UMAP)* plot 10 principal cell types, further confirmed by computational annotation to a reference dataset built from 18 SN samples from an independent study^14^ (***Figure S1E-G)*** and by marker gene expression (***Figure 1B, C***). Representative examples of gene expression uniquely defining a specific cell type include oligodendrocyte transcription factors 1 & 2 (*OLIG1/2)* for oligodendrocyte precursor cells (OPCs), myelin-associated oligodendrocyte basic protein (*MOBP*) for oligodendrocytes (ODCs), the classical astrocytic markers Aquaporin-4 (*AQP4)* and glial fibrillary acidic protein (*GFAP)*, complement and chemokine signaling genes *C3* and *CX3CR1* for microglia, neurotransmitter biosynthetic genes for dopamine (DA) neurons (dopa decarboxylase and tyrosine hydroxylase [*DDC, TH*]) and non-DA neurons representing gabaergic (glutamic acid decarboxylase GAD1/2) and glutamatergic (SLC17A17) neurons, and various markers specific to each of the remaining cell types including endothelium, pericytes, ependyma and T-lymphocytes (***Figure 1B***).

We then asked whether opioid exposure altered the proportions of various cell types. In controls, oligodendrocytes (ODC) and their precursors (OPC) taken together comprised 64.3% of all SN nuclei, a proportion that is highly consistent with an independent dataset^14^, followed by astrocytes (15.0%) and microglia (13.5%). In contrast, DA and GABA/non-DA neurons together accounted for 3.8% of VM nuclei, while pericytes, endothelium, T-cells, and ependyma together represented the remaining 3.1% of nuclei in our VM specimens from control cases. (***Figure 1D***). Of note, our overdose cases showed very similar numbers and proportions for each cell type compared to controls. We conclude that opioid exposure and OD is not associated with proportional shifts among the neuronal and glial constituents in the VM (***FIg. 1D***). Consistent with this observation, case and control groups from each of our two collection areas, when plotted separately into our UMAP coordinates, showed highly similar distributions by cell type (***Fig. 1E***).

### Hundreds of glial transcripts show altered expression in opioid-exposed midbrain

Next, we explored cell-type specific differential gene expression (DEG) by diagnosis (history of opioid use and overdose vs. drug-free control). Focusing on autosomal gene expression, we first obtained a counts matrix of 30,801 genes, presenting in 531 combinations (each comprised of a minimum of 30 single nuclei) defined by sample and cell type. Importantly, the Detroit and Miami cohorts, each analyzed separately, showed on a genome-wide scale strong positive correlations for cell-type specific transcriptome differences between their respective cases and controls, speaking to the generalizability of findings. Specifically, the highest z-score correlations (Miami vs. Detroit) were observed for OPC (R=0.44), ODC (R=0.32), astrocytes (R=0.30) and microglia (R=0.29) (P< 2.2^-10-16^) (***Figure S2A***),

Therefore, in order to further examine cell type-specific gene expression changes in the VM from overdose cases, we next conducted differential gene expression (DEG) analysis by combining the Detroit and Miami cohorts, using sex, genetic ancestry (genotype principal components), age, and postmortem confounders (e.g. brain pH) as covariates. Our initial round of covariate-corrected DEG analysis, with a log fold-change threshold (FCT) >0.25 and FDR corrected P <0.1, we called 5,239 DEGs from a total of 25,728 genes included in the DEG analyses, with 2,999 up- and 2,381 down-regulated, with many of these genes dysregulated in more than one cell type (***Table S3***). However, the impact of OD on the genome-wide VM transcriptome showed striking disparities by cell-type due to the preponderance of glial-specific alterations. Thus, 9.5% (2,131/22,536) of microglia-, and 7.5-6.2% (1,503/20,050 to 1,462/22,536) of astrocyte- and ODC/OPC-associated transcripts were differentially regulated in comparison to drug-free control cases. In sharp contrast, only 0.32% (70/21,881) of the GABA/non-DA neuronal transcriptome, and none (0/15,041) of DA neuron expressed transcripts, were detected as altered in OD cases (***Table S3***). Furthermore, while the largest share of DEG, or 2,131/5,239 (42.5%) was contributed by the microglial population, each of the major glial subtypes shared between 246-395 DEGs with at least one additional glial subtype (***Figure 2***), with shared directionality in a large majority of these DEGs (83-95%, depending on cell type) (***Figure S2B***).

**Figure 2:**
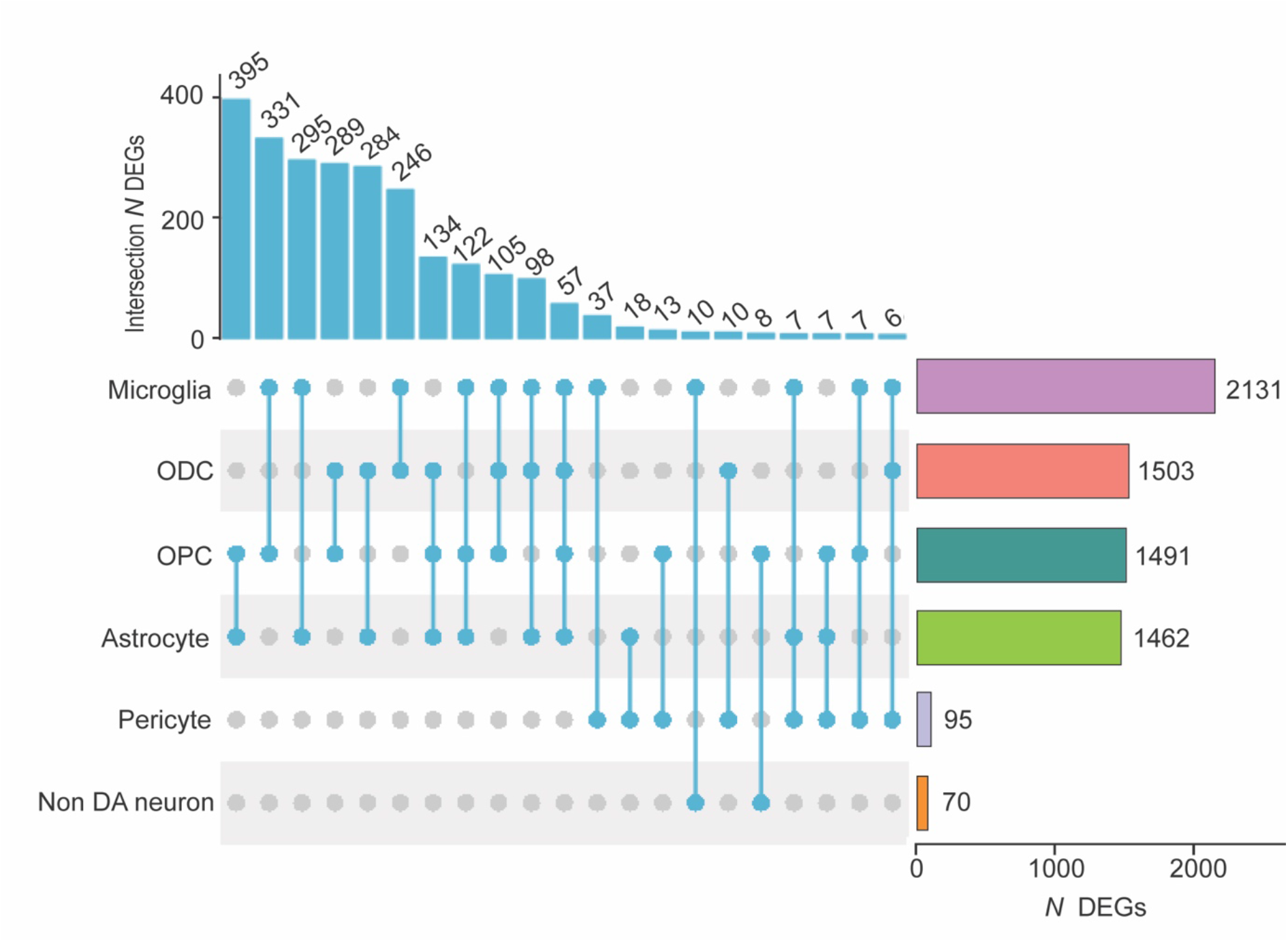
Representation of differentially expressed genes (DEGs) shared across VM cell types. (left) Counts of shared DEGs across cell types, as indicated. (right) Number (*N*) of DEGs for each cell type, from (top) microglia, N = 2131 DEGs to (bottom) non-dopaminergic neurons, N = 70 DEGs.

### Transcriptomic signatures of opioid-exposed midbrain include glial activation and downregulation of synaptic functions in non-dopaminergic neurons

We noted that expression of molecules broadly linked to glial activation and neuroinflammation, including STAT3, STAT5A/B and other members of the STAT (*Signal Transducer and Activator of Transcription*) transcription factor family^15,16^ were upregulated in various glial populations of OD VM (***Table S3***). Therefore, in order to explore this phenomenon on a genome-wide scale, we next conducted cell-type specific gene ontology (GO) over-representation analyses. Indeed, up-regulation of immune response pathways including, for example, interferon, NFkB signaling, and cell motility ranked top in all glial populations of VM, including astrocytes, pericytes, microglia, and ODC/OPC (***Figure 3, Figures S3***,***S4, Table S4)***. In striking contrast to these types of glial activation, top ranking GOs enriched in neuronal DEGs revealed downregulation of functions related to synaptic connectivity including ionotropic glutamate receptor signaling, long-term potentiation, neurite extension and others, in conjunction with increased chromatin repression by histone (H3-lysine 9) methylation (***Figure 3, Figure S4, Table S4)***.

**Figure 3:**
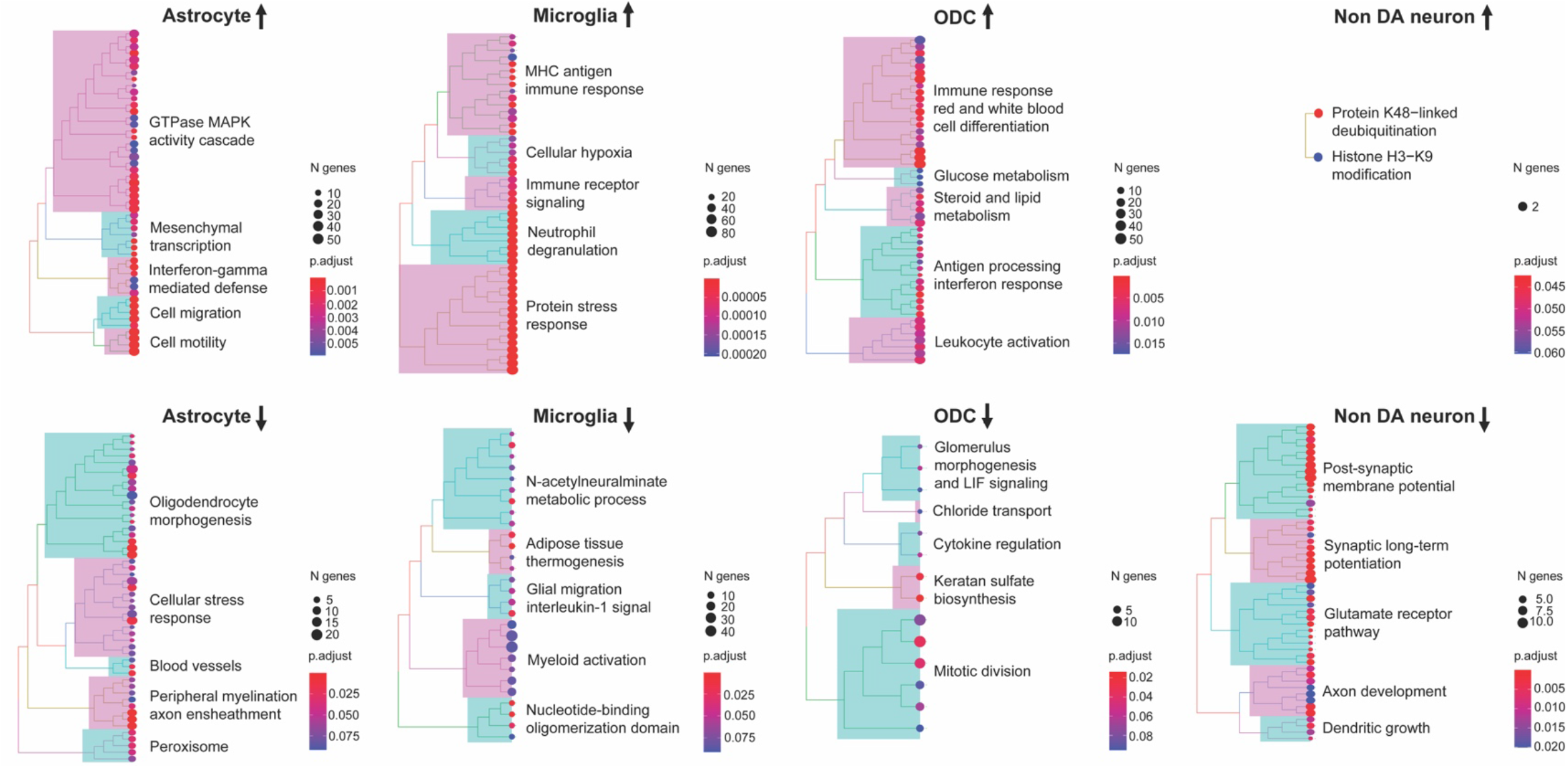
Pathway enrichments of up- and down-regulated VM transcripts by cell type. Biological Process GO enrichments as indicated. For additional details, see Figures S3 and S4 and Table S4.

Having shown widespread reprogramming of the nuclear transcriptome in multiple cell populations of VM from opioid cases, with activation of immune signaling in multiple glial populations and decreased synapse-related gene expression in (non-dopaminergic) neurons, we then asked whether these observations would be broadly reproducible by gene expression profiling from whole cells or even bulk tissue. To this end, we compared the differential gene expression between opioid and control cases in the current study for each VM cell type to an earlier, smaller (N=50) study^10^ involving RNA-seq profiling of bulk VM tissue (NB: there was no overlap in cases used in the two studies). We observed positive z-score correlations that were strongest for the astrocytic (R=0.19), ODC (R=0.18) and microglia (R=0.17) populations (***Figure S5***). These correlations were highly significant (p< 1.96 × 10^−4^). In addition, previously reported GO enrichments from bulk VM tissue analysis strikingly resonated with the findings presented here, including upregulation of NFkB signaling and inflammatory, cytokine and hypoxia response pathways, and were highly reminiscent of the glial activation patterns identified in the present study.

Many individual glial transcripts significantly altered in our opioid cases in a cell-specific manner replicated the most robust changes seen in the aforementioned VM whole-tissue RNA-seq study^10^ (***Figure 4***). For example, among the group of immune signaling genes with increased expression in opioid-exposed VM, up-regulated Interleukin 4 receptor (*IL4R*) expression in VM bulk tissue was previously reported to be strongly predictive of diagnostic categorization (opioid vs control)^10^. According to the cell-specific results presented here, the *IL4R* gene is highly expressed in microglia, consistent with a role in anti-inflammatory reprogramming of microglia and macrophages after brain or nerve injury^17^. Another top ranking gene in the VM opioid overdose whole-tissue RNA-seq study was MAP3K6 kinase, a gene predominantly expressed by astrocytes and implicated in angiogenesis^18^. In the current study, MAP3K6 and the related molecule MAP3K7, implicated in abnormal neurovascular regulation in opioid use disorder and overdose brain^19^, were confirmed as being dysregulated specifically within astrocytic nuclei.. Furthermore, notable pathway alterations in the OPC/ODC cell population of OD cases of the present study included CNS injury-mediated differentiation programs, including the bZIP MAF transcription factor MAFF^20^ and the cell cycle regulator Cyclin Dependent Kinase Inhibitor *CDKN1A* which, again, were among the top scoring DEG in the previous VM bulk tissue-based gene expression study (***Figure 4***). Furthermore, the latter gene is robustly induced in ventral striatum of morphine-exposed mice^21^ and in VM of subjects diagnosed with cocaine use disorder^22^, implicating a broader role for *CDKN1A* in addiction biology beyond opioids in the midbrain.

**Figure 4:**
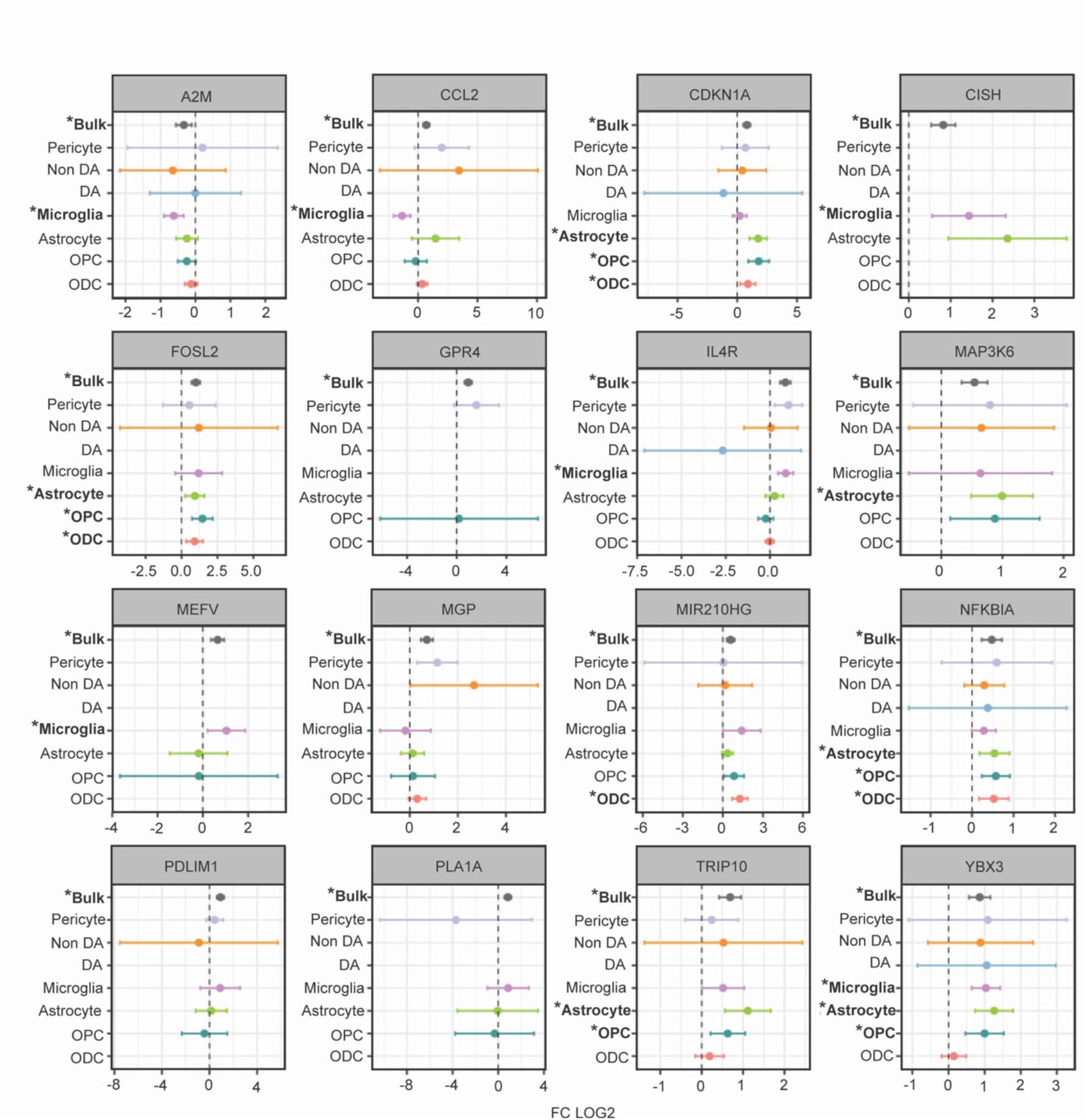
Differentially expressed genes in VM from opioid cases: Comparison of whole-tissue and cell-specific data. Forest plots of selected DEGs (FDR P<0.1) in overdose VM, comparing (x axis log fold change compared to control) previous VM bulk tissue RNA-seq (ref.10) to cell-type specific profilings in current study. *, bold font, FDR P<0.1. See also Figure S5.

We note that in both the previous VM bulk tissue RNA-seq^10^ and the present VM single nuclei RNA-seq studies, neuron-specific transcriptional programs for structural and functional neuronal connectivity, and synaptic transmission were downregulated in the opioid group. According to the present study, this effect is driven by transcriptomic alterations in the non-dopaminergic neurons. Furthermore, the AP-1 transcription factor and early response gene, FOSL2, previously found to be induced in rodent addiction circuitry by chronic morphine administration^23^, showed increased expression in numerous neuronal and glial cell populations of our sn-RNA-seq study, including OPC, microglia and non-dopaminergic neurons. Similarly, FOSL2 was among the top scoring DEGs in the bulk VM RNA-seq study (***Figure 4***).

To summarize, the current study identified robust changes in human VM gene expression and associated GO pathways in cases with a history of opioid use and OD that are consonant with a previous report, but reveal the cell type-specific nature of these changes, which can be characterized as broad glial activation with a prominent representation of immune signaling pathways, in conjunction with downregulation of glial support and neuronal signaling genes. This transcriptomic signature observed was independent of the period of specimen collection or geographical location of the cohorts.

### Cell-specific DEGs associated with genetic risk for Addiction Disorder

Next, we wanted to explore whether any cell type-specific DEGs in our opioid exposed cases are associated with the genetic risk for addiction disorder. Of note, while opioid use disorder is considered moderately heritable, with an estimated 60% of population variability attributable to genetic factors^24,25^, to date only two or three loci have been genome-wide reproducibly linked to opioid use and substance-associated traits^26^. Remarkably, these loci include the cell motility regulator, *SCAI* (chr. 9q33.3)^26^, which in our study is significantly down-regulated in VM microglia from opioid cases (***Table S3***). However, opioid exposure is broadly associated with genetic risk for substance use and dependence overall^27^. Therefore, we wanted to explore whether any gene expression alterations in our opioid cases, including cell type-specific dysregulation, could match a broader list of genes linked to heritable substance use traits. To this end, we screened PhenomeXcan^28^, a resource for transcriptome-wide association studies linking genes to phenotypes by genetically predicted variation in gene expression. We focused on brain gene expression and population-scale substance use phenotypes in PhenomeXcan including 40 traits related to caffeine, nicotine, alcohol, and marijuana consumption or dependence^29^ (***Table S5***) and, for comparison, a number of medical traits associated with hundreds or thousands of significant genes in the PhenomeXcan resource. While we found no specific enrichment of our cell type-specific DEG datasets with traits linked to any specific drug or substance (***Figure S6***), among the 1,260 PhenomeXcan genes linked to a substance use trait and expressed by at least one cell type in the present study, 325, or 25.8%, were called as significantly altered in our study, including 149 DEGs in microglial nuclei, and 100-70 DEGs in astrocytes and oligodendrocytes and their precursors, respectively, and only very minimal contributions from some of the remaining cell types including pericytes and non-dopaminergic neurons (***Figure S7, Tables S6, S7***). This included 17 genes called as DEG in one or more VM cell types of the present study, and in the VM bulk tissue RNA-seq study^10^, with 14/17 of genes showing the same direction of change across studies (***Table S5***). Among these, *NUPR1*, a regulator of chromatin acetylation, showed upregulation of expression in VM tissue^10^ and VM astrocytes of our opioid cases, and was identified as a risk gene in caffeine, alcohol and marijuana abuse and dependence Interestingly, NUPR1, also known as *STRESS PROTEIN 8* or p8, sensitizes astrocytes to oxidative stress when upregulated^30^, in effect exerting a protective effect by decreasing the production of oxygen radicals^31^. Other notable SUD PhenomeXcan genes up-regulated in VM astrocytes, and in VM tissue^10^, of opioid cases include the A2B adenosine receptor (*ADORA2B*), which regulates synaptogenesis and synaptic plasticity by downregulating glutamate receptor 5 signaling in astrocytes^32^. Furthermore, the *NFKB2* transcription factor, previously linked to tobacco smoking by transcriptome-wide association, was upregulated in our study in multiple glial subtypes including microglia, astrocytes and OPC of our opioid cases, again consistent with previous observations made in VM tissue^10^. Interestingly, *NFKB2*, a broad regulator of the genomic response to inflammatory stimuli, is down-regulated in the brains of rats after extinction training for cocaine self-administration^33^ in a model for cocaine use disorder. Furthermore, the *Cytokine Inducible SH2 containing Protein (CISH)*, linked to substance use in PhenomeXcan, which was upregulated in VM microglia of our overdose cases, and in the VM bulk tissue RNA-seq study^10^, encodes a JAK/STAT pathway signaling molecule linked to the anti-inflammatory response in that cell type^34^.

In addition to this set of 17 substance use-associated genes that showed significant expression changes in the opioid group, we also identify a new set of 312 PhenomeX substance use-associated genes with significant expression changes in multiple glial subtypes of opioid-exposed VM and including several genes linked to the pharmacogenomics of opiate use disorder. For example, *GSG1L* has been linked to plasma methadone levels^35^, and the myeloid zinc finger transcription factor MZF1 to cis-regulatory sequences driving expression of the mu-opioid receptor 1^36^. Another noteworthy example of a PhenomeX substance use gene with altered expression in all three major glial prototypes, including astrocytes, microglia and ODC/OPC (***Figure 5***) is the nuclear paraspeckles-associated long non-coding RNA, *NEAT1*, which has been broadly linked to astrocytic and microglial activation^37,38^, and has been previous reported as up-regulated in ventral striatum of heroin abusers^39^. Consistent with these findings, multiple inflammatory response- and cell activation- and migration-associated GOs involving *NEAT1*, and additional PhenomeX substance use-associated glial DEGs such as *ATP1B3, BARD1* and *HESX1* (***Figure 5***) were significantly enriched among the glial DEGs of the present study (***Table S4***).

**Figure 5:**
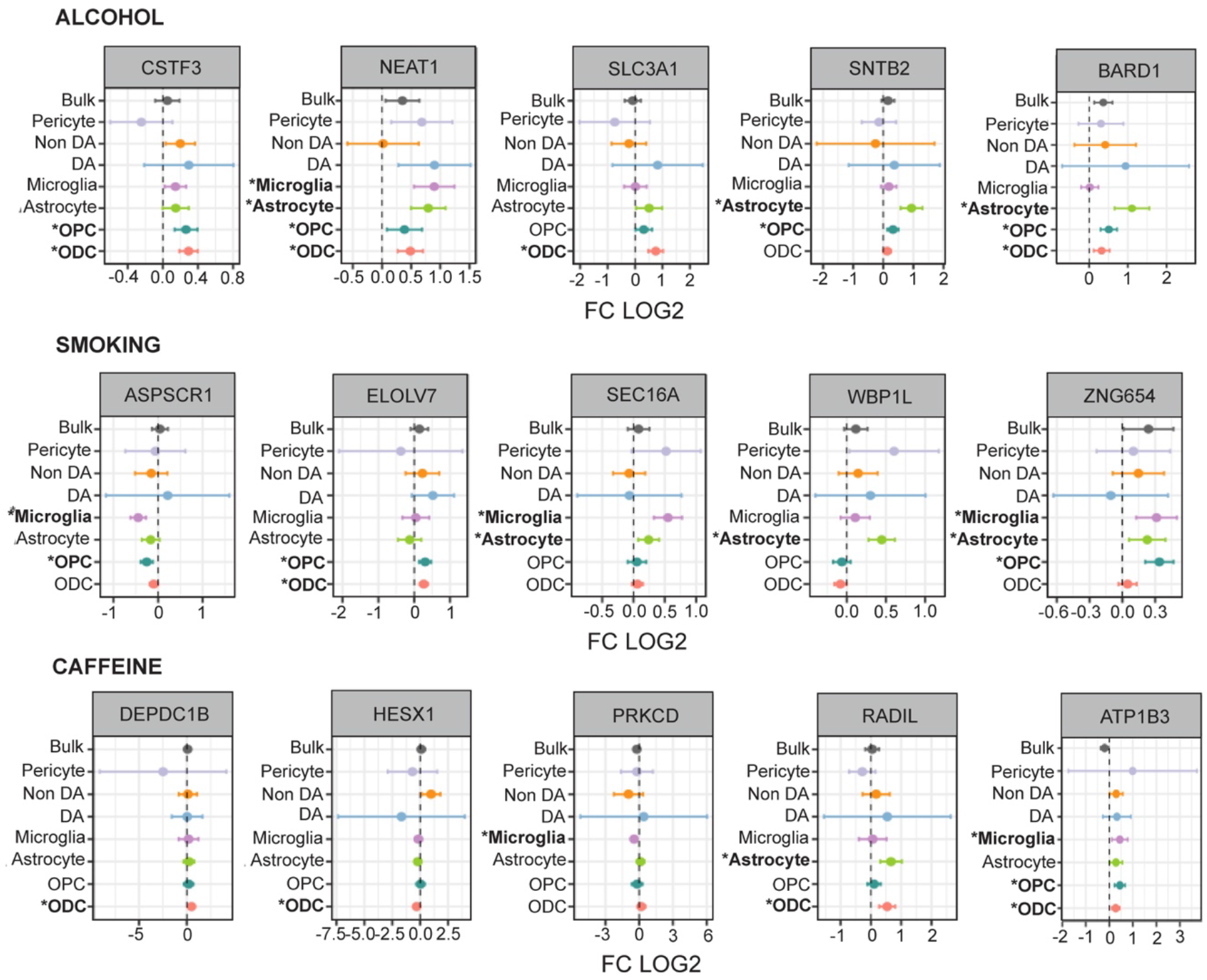
Differentially expressed genes in the VM of opioid cases linked to substance use in the general human population. Forest plots of selected DEGs (FDR P<0.1) in OUDd VM (x axis log fold change compared to contro; y-axis VM bulk tissue and single nuclei by cell-type RNA-seq) that are linked to alcohol and caffeine consumption, and smoking in PhenomeX database. *, bold font, FDR P<0.1. See also Tables S4, S5.

## DISCUSSION

### The glial response to opioid exposure

The present study, which involved cases with a history of opioid use and dying of opioid overdose matched with drug-free controls, from two independent specimen collection sites in geographically distinct areas of the continental U.S., is one of the largest postmortem studies in this field, profiling the transcriptome in a total of 212,713 single nuclei from the ventral midbrain. Key findings included up-regulation of pro-inflammatory cytokine and immune response pathways, NFkB signaling and generalized activation affecting all major glial populations including microglia, astrocytes, and oligodendrocytes and their precursors. In contrast, transcript reprogramming in VM neurons, and specifically in the non-dopaminergic (and thus mostly GABAergic) neuronal population, was defined by down-regulated expression of genes involved in synaptic plasticity and neuronal connectivity. Furthermore, there was significant overlap in differentially expressed genes and associated Gene Ontology pathways between the current VM single nuclei RNA-seq study, and a previous VM bulk (whole-tissue) RNA-seq study^10^ of a smaller but distinct cohort of subjects, which showed a similar activation of inflammatory signaling. Thus, the current study identified robust changes in human VM gene expression and associated GO pathways in cases with a history of opioid use and overdose that are consonant with the previous report, but revealed the cell type-specific nature of these changes, which can be characterized as broad glial activation with a prominent representation of immune signaling pathways, in conjunction with downregulation of glial support and neuronal signaling. It is remarkable that the aforementioned transcriptomic signatures were independent of the period of specimen collection and geographical location of the cohorts.

Recent transcriptome profiling of bulk tissue from prefrontal cortex and nucleus accumbens (using an independent brain collection) reported prominent upregulation of immune signaling genes and transcriptomic signatures indicative of microglial activation and glial motility in the forebrain associated with opioid exposure and overdose^40^. Moreover, a RNA-seq study in ventral striatum of mice exposed to morphine, conducted on FACS-sorted oligodendrocytes in conjunction with a general survey of striatal transcription at single nucleus resolution, also broadly resonate with the current findings by demonstrating drug-induced activation of stress and differentiation pathways in all major glial prototypes, with some key genes (e.g. *CDKN1A* and *NFKBIA)* found differentially regulated in the ODC and astrocytes of opioid VM samples in the present study also changed in the opioid-exposed mouse^21^(***Table S8***). For a subset of differentially regulated glial activation markers, including Glial Fibrillary Acidic Protein (GFAP), opioid-induced up-regulation in VM was first reported, in a rat model, three decades ago^41^.

These studies, taken together, lead to a remarkably consistent consensus implicating glial activation with up-regulation of inflammatory and immune signaling pathways across multiple nodes in the addiction circuitry of the opioid exposed brain, including the ventral midbrain. Therefore, it is extremely interesting that drugs acting as inhibitors for the pro-inflammatory glial response reduce morphine-induced withdrawal effects in the rat model^42^, and attenuate the addictive features of opioids, including positive reward (for example, ‘feeling high’) and withdrawal-associate symptoms, in human volunteers diagnosed with Opioid Use Disorder^43–45^. The precise pharmaco-molecular and -cellular cascades linking opioid addiction and dependence to glial activation remain to be elucidated. Potential mechanisms could include drug-induced activation of TLR4 and other *Toll-like* receptors which, in turn, activate NFkB to drive transcriptional activation of cytokine and chemokine^46^. Furthermore, there is evidence for functional interaction between opioid-receptor and NFkB signaling as discussed in^40^. Given that the transcriptomic alterations in striatum of subjects with cocaine use disorder point to an opposite effect, i.e., decreased neuroinflammation^47^, interventions against the pro-inflammatory effects of opioids could provide unique drug class-specific therapeutic opportunities.

### Limitations of the present study

The present study provided new insights into cell type-specific gene expression changes in each of the major glial populations, including OPC, ODC, astrocytes, and microglia, within the VM of opioid overdose cases. However, our experimental design included pooling of up to 4 VM specimens for 10K nuclei target recovery for each 10x chromium gel bed assay and, in agreement with previous single nucleus RNA-seq work on human VM^12-14^, less abundant cell types such as dopaminergic and non-dopaminergic VM neurons, as well as pericytes, T-cells, and endothelial and ependymal cells, each comprise only a very small fraction (0.8% - 3%) of the total population of VM nuclei in the current study. Multiple case and control samples lacked the minimum number of nulclei from some of these cell types in order to enter DEG analysis by cell type (set at N ≥ 30 nuclei per cell type, see methods), reducing the overall power for many of these rarer VM cell populations (***Table S2***). Future studies that specifically enrich for cell types such as VM dopaminergic neuron nuclei based on fluorescence-activated nuclei sorting^48^ will be required in order to more fully elucidate opioid-induced transcriptional alterations for these relatively rare VM cell types.

### Neuron-specific alterations in VM from opioid overdose cases

The present study identified 69 genes dysregulated in the non-dopaminergic neuronal population in the opioid exposed VM (***Table S3***). These included neuropsychiatric risk genes such as *AUTS2* and *NCALD*, which reportedly are transcriptionally dysregulated in a VM target, the ventral striatum, after cocaine and amphetamine abuse^49,50^. Furthermore, the aforementioned AP-1 transcription factor *FOS*L2 was up-regulated in VM non-dopaminergic neurons. We counted 9/69 (13%) of differentially regulated neuron-specific genes in VM in our opioid group that matched to neuron-specific enhancer or promoter sequences reportedly affected by histone hypo-acetylation in prefrontal cortex (PFC) neurons from opioid overdose cases^51^ (***Table S9***). These included genes conferring heritable risk for nicotine and other substance dependence, such as *GABBR2* encoding a GABA_B_ receptor, and *SHC3* involved in MAP kinase and neurotrophin signaling ^52,53^. These findings then further support the emerging hypothesis that opioid exposure and addiction is associated with a coordinated transcriptional dysregulation in various neuronal and non-neuronal subpopulations residing in specific nodes of addiction circuitry including VM, striatum, and PFC^54,55^.

We predict that future studies, by constructing a neurogenomic atlas of cell specific gene expression alterations and related changes in epigenomic regulation, across multiple regions of opioid-exposed brains and conducting integrative analyses combined with the emerging genetic risk architecture for substance use disorders, will provide deep insight into neuronal and glial mechanisms highly relevant to the neurobiology and treatment of opiate addiction.

## FIGURE LEGENDS

**Supplementary Figure 1: Quality controls for VM single nuclei RNA-seq profiling. (A)** Dexmulet genotyping confirms 100% match by donor by pool against the background of all 95 donors. Y-axis represents the library pools for 95 individuals. X axis represents the individual, prefix of which denoting the expected library for each individual. Heat bar, percentage of nuclei for specific a individual in library, as indicated. **(B-D)** Violin plots, showing separately for 45 opioid cases (blue) and 50 controls (red), **(B)** the distribution of numbers of nuclei per sample, **(C)** the numbers of genes per nucleus, and (**D**) numbers of reads per sample. (**E**) UMAP plot for VM from 95 samples (present) study, (**F, G**) automatic annotation of major VM cell types from previous study in small reference sample^14^.

**Supplementary Figure 2**: **Differential gene expression across cohorts and cell types**. (**A**) Z-score (case control differential) correlations comparing (y-axis, Miami; x-axis Detroit) cohorts for 4 major glial populations, as indicated, and non-dopaminergic neurons in the VM. (**B**) Proportional representation of DEGs shared between specific pairs of glial subtypes, as indicated on x-axis, based on shared vs. opposite directionality.

**Supplementary Figure 3: Cell-type specific pathway enrichments of differentially expressed genes in astrocytes and microglia**. GO pathway enrichments for Biological Process, shown separately for up- and down-regulated DEGs, as indicated. See also Figure 3 and Table S4.

**Supplementary Figure 4: Cell-type specific pathway enrichments of differentially expressed genes in oligodendrocytes (ODC) and non-dopaminergic neurons (NON-DA)**. GO pathway enrichments for Biological Process, shown separately for up- and down-regulated, as indicated. See also Figure 3 and Table S4.

**Supplementary Figure 5**: **Correlations between prior bulk RNA-seq study and the current RNA-seq analysis with single cell resolution**. Z-score (case control differential) correlations comparing (y-axis, bulk tissue RNA-seq study^10^; x-axis, cell type specific RNA-seq at single nuclei resolution, current study) for 7 cell types, as indicated.

**Supplementary Figure 6**: **DEGs linked to TWAS**. Percentages representing, for each cell type the normalized proportion of DEGs overlapping with PhenomeXcan TWAS for substance use and medical or neurological traits, as indicated.

**Supplementary Figure 7: TWAS DEG by cell type**. Cell-type specific counts of genes called as DEG in current study *and* linked to substance use trait(s) in PhenomeXcan (see Table S6).

## FUNDING

This work was supported by NIH National Institute of Drug Abuse R01 DA047880.

## DATA AND CODE AVAILABILITY

Raw and processed data is in the process of being submitted to dbGAP/GEO/SRA. Original code and scripts used to analyze the data will be made available on GitHub.

## ACKNOWLEDGEMENT

We thank the Icahn School of Medicine at Mount Sinai’s Flow Cytometry Core for providing expertise and support on nuclei sorting, the Scientific Computing group at the Icahn School of Medicine at Mount Sinai for computational resources, and personnel at the New York Genome Center for sequencing support.

## METHODS

Case and control brains were collected within two separate geographical areas of the U.S., representing the greater Detroit and Miami metropolitan areas. The two brain specimen repositories operate independently.

### Detroit

Human midbrain specimens were collected during routine autopsy and de-identified specimens were characterized as described previously^10,22,56^. Briefly, cause of death was determined by forensic pathologists following medico-legal investigations evaluating the circumstances of death including medical records, police reports, autopsy results, and toxicological data. Case inclusion in the opioid abuse group was based on a documented history of opioid abuse, toxicology report positive for opioids, and forensic determination of opioids as cause of death. Opioid abusers with a positive toxicology for common non-opioid drugs of abuse (e.g. alcohol, cocaine, cannabinoids, anxiolytics, barbiturates) were excluded from the study, with the exception of inclusion of a subset of cases positive for opioids plus benzodiazepines, as this reflects a drug class commonly co-abused with opioids with deadly consequences^57^. Cases in the control group had no documented history of drug abuse, and tested negative for opiates, cocaine, and other drugs of abuse or CNS medications at time of death. Causes of death for control cases were primarily cardiovascular events or gunshot wounds. Exclusion criteria for either group included a known history of neurological or psychiatric disorder, death by suicide, evidence of neuropathology at autopsy, debilitating chronic illness, estimated postmortem interval > 20h, or biochemical evidence of poor tissue sample quality or prolonged perimortem agonal state (i.e. brain pH <6.2 or RNA integrity number [RIN] <6.0). To reduce variance unrelated to drug abuse, the two groups were matched in terms of gender, race, age, brain pH, and RIN at the time of processing. Table S1 includes demographic and sample quality characteristics. Brains were sectioned transversely at the level of the posterior diencephalon and mid-pons to obtain a tissue block encompassing the entire human midbrain (corresponding approximately to plates 51–56 of DeArmond et al, 1989^58^.

### Miami

Cases were selected from an opportunistic sample of opioid intoxication deaths defined by circumstances of death and forensic and supplemental toxicology data. All cases and unaffected controls were evaluated to rule out comorbid psychopathological diagnoses. Common drugs of abuse and alcohol and positive urine screens were confirmed by quantitative analysis of blood and brain. Retrospective chart reviews were conducted to confirm history of opioid abuse, methadone or addiction treatment, drug-related arrests or drug paraphernalia found at the scene. Supplemental brain toxicology was done on select cases for comparison to blood levels at the time of death. Case inclusion in the opioid group was based on a documented history of opioid abuse, toxicology report positive for opioids, and forensic determination of opioids as cause of death. The detection of 6-acetyl morphine (6-AM) was taken as definitive evidence of acute heroin exposure. Cases of polysubstance abuse, where there was positive toxicology and a history of use of cocaine, methamphetamine, alcohol, and other drugs, were excluded from the study. Drug-free control subjects, with negative urine screens for all common drugs and no history of licit or illicit drug use prior to death, and with no known history of neurological or psychiatric disease, were selected from accidental (motor vehicle accidents or trauma) or cardiac sudden deaths. All cases and unaffected controls were selected from persons who died suddenly without a prolonged agonal state, since agonal state affects brain tissue quality control metrics. Care was taken for cohort selection to match subject groups as closely as possible for non-Hispanic Caucasian ancestry, age, gender, postmortem interval (PMI), RNA integrity number (RIN) values, and brain pH (Table S1).

#### Sample processing

Sample processing for both brain cohorts, including purification of nuclei, RNA extraction, and generation of single nuclei RNA-seq libraries, was performed in New York. Brains were processed in pools of N=3-4 unique brains (donors). From each unique brain, a tissue aliquot, containing approximately 20mg of ventral midbrain from the area of the substantia nigra (A10) and portions of the adjacent ventral tegmental area (A9), was homogenized using a douncer at least 20x in 1ml lysis buffer (0.32M sucrose, 5mM CaCl2, 3mM Mg(Ace)2, 0.1mM EDTA, 10mM Tris pH8, 0.5mM DTT, 0.1% Triton X-100) with 400U RNase inhibitor (Takara Bio Recombinant RNase Inhibitor, Cat. 2313) added to it. Then, an additional 4ml of lysis buffer was added and the sample solution was dounced an additional 20x until homogenous. After douncing, each pool of 4 unique samples was transferred to an ultracentrifuge tube (Beckman Coulter Polypropylene Centrifuge Tubes ⅝ x 3 ¾ in., Ref. 361707) and underlayed with 9ml of sucrose buffer (1.8M sucrose, 3mM Mg(Ace)2, 0.5mM DTT, 10mM Tris-HCl pH8), then ultracentrifuged with 24,000rpm in a SureSpin 630 (17mL) Rotor (106,803x g) for 1hr at 4°C. After centrifugation, the supernatant was removed and each pellet of nuclei was carefully resuspended in 1ml of 1% BSA with 1,000U RRI added to it, transferred to a sterile tube, and 1 µl of the nucleophilic dye, DAPI (4’,6-Diamidino-2-Phenylindole, Dihydrochloride, Invitrogen Cat. D1306), was added. For FACS collection, sterile tubes were coated with 5% BSA. After residual BSA solution at bottom of tubes was removed, DAPI+ nuclei were sorted into the collection tubes using a BD FACSAria Cell Sorter, with approximately 300,000 DAPI+ nuclei for each pool of 4 unique midbrain samples collected and processed using the 10x Chromium Next GEM Single Cell 3’ v3.1 (Dual Index) Protocol (CG000315 Rev A) according to the manufacturer’s instructions. The Agilent 2100 High Sensitivity DNA Bioanalyzer Kit was used as a quality control step at Step 2.4 and end of the library preparation, also as per 10x Genomics’ guidelines. To prepare samples for sequencing, sample concentration was determined using the KAPA Biosystems Library Quantification Kit (ROX Low qPCR Master Mix, Cat. KK4873).

Libraries were sequenced by the New York Genome Center using the Illumina NovaSeq platform aimed at a sequencing depth of 50,000 read pairs per nucleus. Libraries consisted of paired-end reads with a read length of 100bp.

#### Genotyping

Miami Cohort: Genomic (g) DNA was extracted from cerebellar tissue using a QIAamp DNA Micro Kit, followed by SNP genotyping was done with Illumina’s MEGA multiethnic array at Rutgers University Cell and DNA Repository (RUCDR Infinite BiologiX).

Detroit Cohort: DNA was extracted from 25mg of tissue using QIAamp DNA mini kit from Qiagen Cat# 51304 (Qiagen, Germantown, MD). Tissues were lysed manually and then processed through the QIAcube DNA isolation protocol. For genotyping Infinium Global Diversity Array-8 v1.0 Kit microarrays were processed by the Advanced Genomics Core of University of Michigan (Ann Arbor, MI, USA). Genotype information was converted to vcf format using “iaap-cli gencall” and “gtc_to_vcf.py” from Illumina.

For both cohorts imputation was performed using the TOPMed Imputation Server version 1.5.7 (https://imputation.biodatacatalyst.nhlbi.nih.gov) to a total of 292,140,970 genetic variants. The vcf files from the two cohorts were then merged and filtered for high-quality imputation and coverage for at least ten scRNAseq transcripts using bcftools resulting in a vcf file with 8,184,813 genetic variants.

### scRNA-seq raw data processing (Alignment and demultiplexing)

We processed each scRNAseq library using ***cellranger (V7*.*0) count*** using the GRCh38 human reference genome and the default parameters except for the --include-introns option which is recommended for single nuclei preparations. The resulting aligned reads bam files were further processed with ***demuxlet***^*59*^ together with the genotype vcf files. Demuxlet assigns the most likely individual for each cell barcode based on the reads overlapping genetic variants. Based on the demuxlet assignments, we removed barcodes that were called doublets, ambiguous, or that could not be assigned to an individual that corresponded to the pool that was used to create the library. Any library that did not contain at least a case and a control individual was also excluded. Additionally we removed any cells with mitochondrial read rates higher than 20%. These filtering procedures resulted in a total of 212,713 high-quality cells with 36,601 genes across 95 individuals. A median value of 2,008 cells are detected for each sample, with a median value of 8,274 reads per cell and 3,070 genes per cell (***Figure S1B-D***).

### Clustering analysis and cell type annotation

We employed the standard pipeline of ***Seurat R package (v4*.*0)*** to process our scRNA data^60^. We first utilized the log1pCP10K approach to normalize data and then standardized the gene expression across cells. For dimensionality reduction analysis, we performed principal component analysis (PCA) on 2,000 highly variable genes to obtain 100 principal components (PCs). To correct batch effects, we calculated the harmony-adjusted PCs using ***RunHarmony*** considering various library effects^61^. The Uniform Manifold Approximation and Projection was used to visualize our scRNA data using the top 50 harmony-adjusted PCs. For cluster analysis with the top 50 harmony-adjusted PCs, we identified 14 distinct clusters across samples at 0.07 resolution.

To annotate cell-type for the scRNA data, we performed differential gene expression analysis between the compared cluster and the remaining clusters (grouped together) to identify cluster-specific expressed genes. The canonical cell-type marker genes were expected to be highly expressed in the corresponding clusters (***Figure S1E-F, Figure 1B***): (1) Oligodendrocytes (ODC) marker genes (such as *MOBP, MBP, PLP1* and *CNP*) were highly expressed in clusters 0, 5, 10 and 13; (2). astrocyte-marker genes (such as *GFAP, AQP4* and *SLC1A2*) are mainly enriched in the cluster 1; (3) microglia-marker genes (such as *C3, CSF1R, CX3CR1, LRRK1, DOCK8* and *P2BY12*) are highly expressed the cluster 2 and 11; (4) oligodendrocyte precursor cells (OPC)-marker genes including *VCAN, PDGFRA, OLIG1 and OLIG2* are mostly expressed in the cluster 3; (5) dopaminergic neuron (DaN)-specific genes such as *DDC, SLC6A3, SLC18A2*, and *SLC18A2* tend to be highly expressed in the cluster 7; (6) Non-dopaminergic (Non-DA) neuron genes including *GAD1, GAD2* and *SLC17A7* gene family tend to be highly enriched in the cluster 4; (7) pericytes-marker genes (*MYO1B, NR4A2, PDGFRB* and *RGS5*) are highly expressed in the cluster 6; (8) While endothelial-specific genes such as (*CDH5, CLDN5, FLT1, KDR, PECAM1* and *PTPRB*) are highly expressed in cluster 8; (8) We also identified some T cell specific genes including *CD96, IL7R, SKAP1* and *THEMIS* highly expressed in the cluster 9; (9) For the cluster 12 with very few number of cells, the genes including *HTR2C* and *TTR* are highly enriched, which are related to the function of ependymal cell.

We also inspected our dataset with an automatically generated cell-type annotation using an independent high quality reference data set^14^ collected from human substantia nigra (SN). This reference set is comprised of 387,483 cells across 18 samples, annotated by 7 major cell types, including ODC, astrocyte, microglia, OPC, dopamine neurons (DA), Non_DA neurons and endothelial^14^. We constructed a heatmap to visualize in our dataset the proportion of different cell types accounting for each cluster (***Figure S1G***). We note that the above 7 major cell types are dominant in the corresponding clusters that highly expressed cell-type marker genes. After combining the cell type marker genes and automatic cell-type annotation, eventually we assigned the cluster 0, 5, 10 and 13 to be ODC, the cluster 1 as astrocytes, cluster 2 as microglia cells, cluster 3 as OPC, cluster 4 as Non-DA neuron, cluster 6 as pericytes, cluster 7 as DaN, cluster 8 as endothelial cells, cluster 9 as T-cells and cluster 12 as ependymal-cells. This cell-type annotation was used for all downstream analyses.

### Differential gene expression analysis

We generated the pseudo-bulk counts data by summing the reads for each gene across cells that were from the same sample and cell-type. Focusing on the autosomal genes, we obtained a counts matrix of 30,801 genes in 531 combinations of the sample and cell-types with at least 30 nuclei (in pilot studies, we varied the minimally required number of nuclei per cell type and sample from 20 to 100, and determined N = 30 nuclei per sample and cell types as minimum number to enter into the case vs. control DEG because smaller N’s increased overall noise factor in the DEG and higher N’s were overly restrictive by excluding larger number of subjects for some of the rare cell type). Combinations were also eliminated if the remaining batches did not include at least one individual from both the control and opioid group. This resulted in 7 cell-types considered for the final differential gene expression analysis. For four of these cell-types we have a large number of individuals, while for three the number of individuals is more reduced and the statistical power is more limited. We performed differential gene expression analysis between opioid and control groups for each cell-type separately using R ***DESeq2 package***^*62*^ and the following model

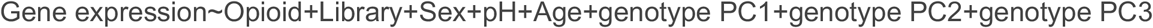

which would correct the noises arising from various experimental batch, ethnicity and other covariates (gender, age and pH). To filter out lowly expressed genes, we only considered genes that were expressed higher than 0.5 CPM in at least 3 samples from opioid and control (respectively) across libraries. To correct for multiple hypothesis testing, we used the Benjamini–Hochberg approach implemented in ***results*** function in the ***DESeq2 package*** with default parameters. We defined the differentially expressed genes as those with False Discovery Rate (FDR) < 10% and the fold change at least 1.189.

### Gene Ontology (GO) enrichment analysis

Using the ***ClusterProfiler (4*.*0) R package***^*63*^, we performed Gene Ontology (GO) enrichment analysis for DEGs from the 6 cell types for up-regulated and down-regulated genes separately. To correct for multiple hypothesis testing, we applied the Benjamini–Hochberg approach to calculate the FDR across all the 12 conditions (6 cell types × 2 directions) implemented in ***p*.*adjust*** in R (4.1). The significantly enriched biological process (BP) terms are defined as those terms (gene size ranging from 5 to 500) with FDR<10%.

### Transcriptome-wide association analysis (TWAS)

To investigate whether DEGs are associated with the genetic risk variants for substance use traits and disorder (SUD), we screened the ***PhonomeXcan database***, a resource for transcriptome-wide association studies linking genes to phenotypes by genetically predicted variation in gene expression^28^. We focused on Substantia Nigra (SN) gene expression and population-scale SUD phenotypes in PhenomeXcan. We focused on a total of 40 SUD-related traits from the following categories of traits or diseases: addiction, alcohol, caffeine, marijuana, and smoking (***Table S5***). We conducted the proportion test using ***prop*.*test*** in R(4.1) to examine whether the DEGs overlapping with SUD are enriched in some cell type or some specific trait. We also integrated other polygenic traits (including BMI, height, CAD, ADHD and Parkinson’s Disease). In the trait-specific analysis, we compared to those from all the traits of interest while those across cell types being the contrasted group in the cell type-specific analysis.

## Notes

Conflicts: The Authors report no conflicts of interest.

Funding: Supported by NIH grant R01DA047880.

### Competing Interest Statement

The authors have declared no competing interest.

