## Supplementary figures and images for "Single Nucleus Transcriptomics Reveals Pervasive Glial Activation in Opioid Overdose Cases"

### Supplementary Figures S1-S7

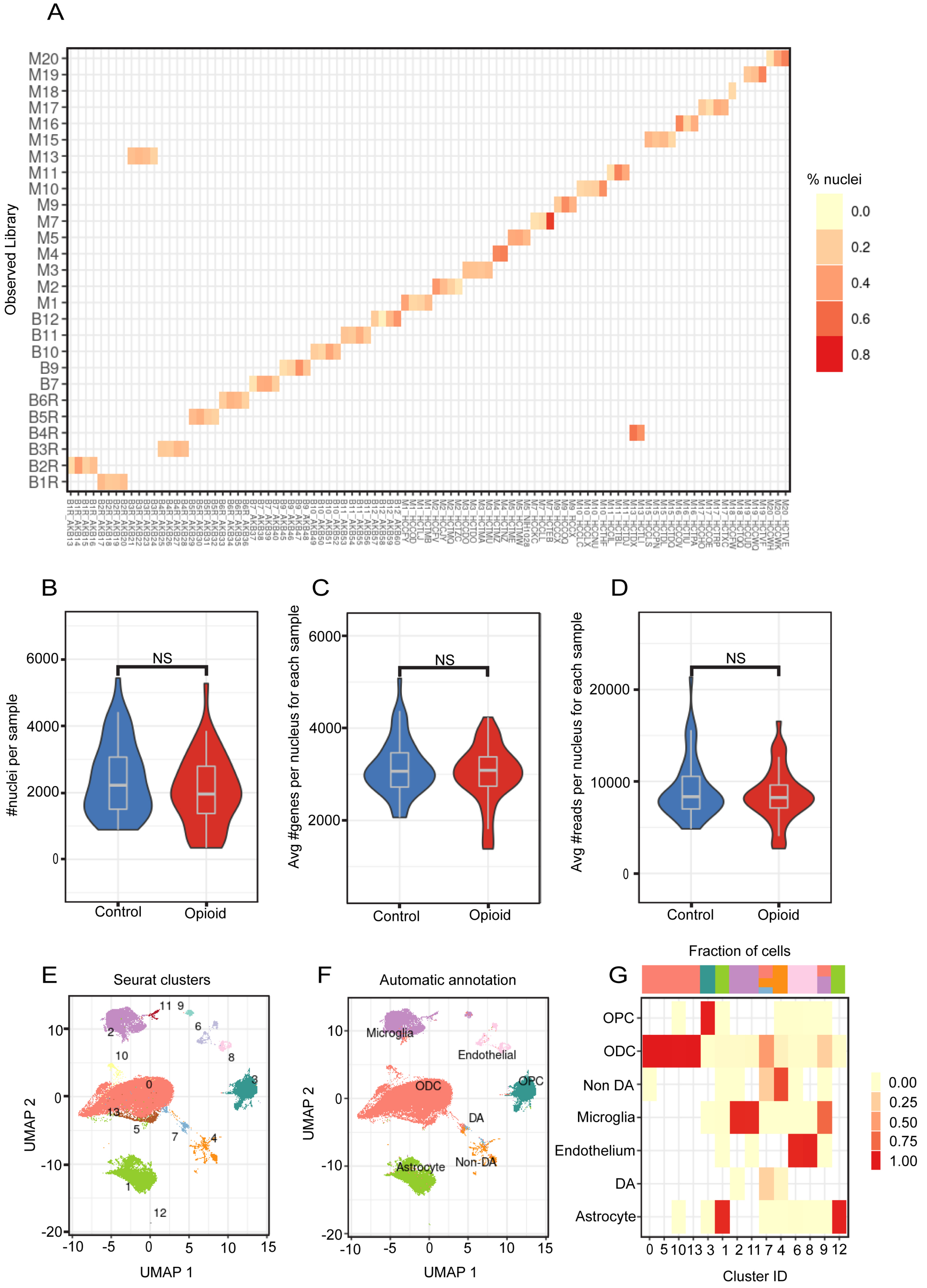

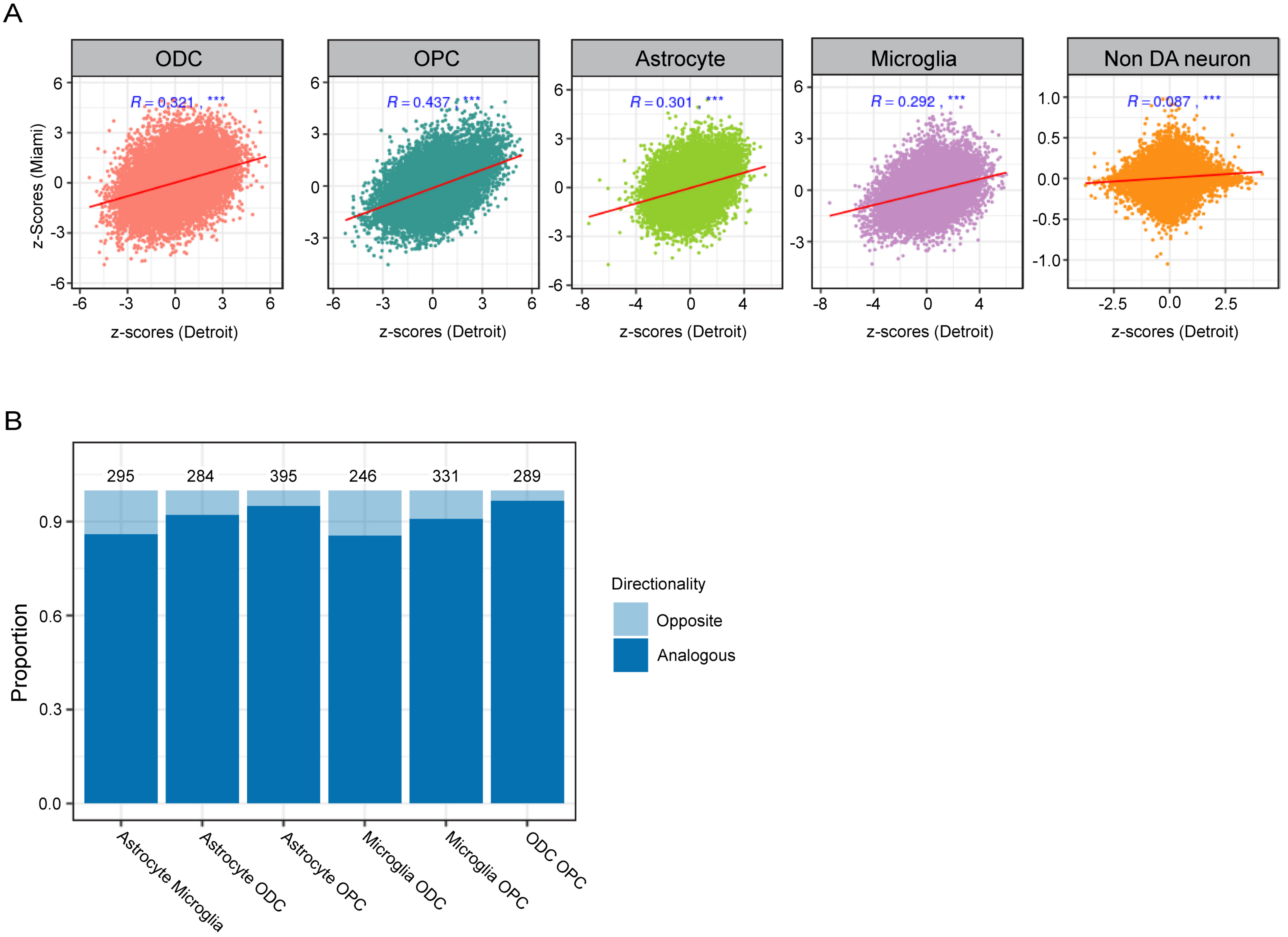

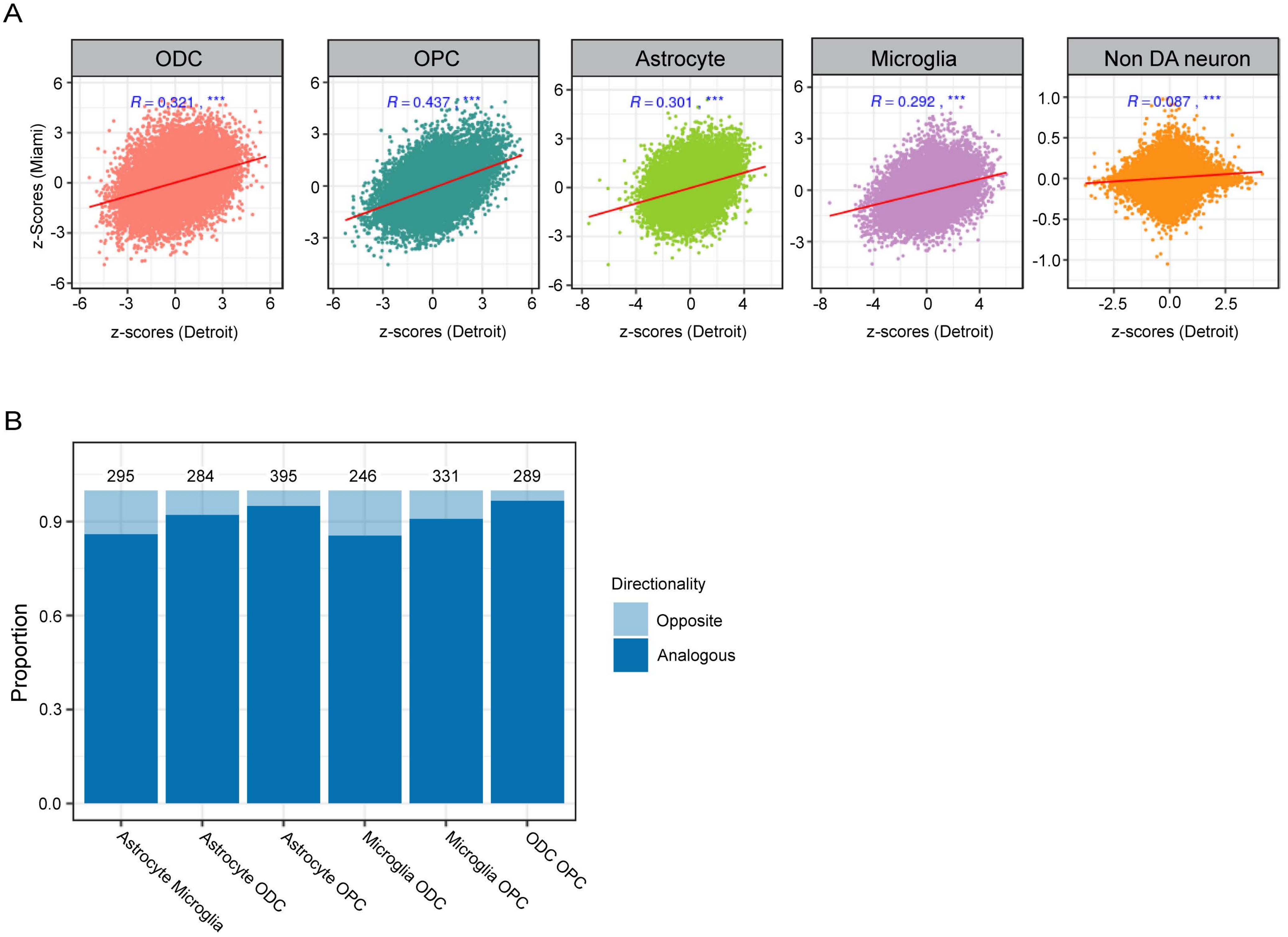

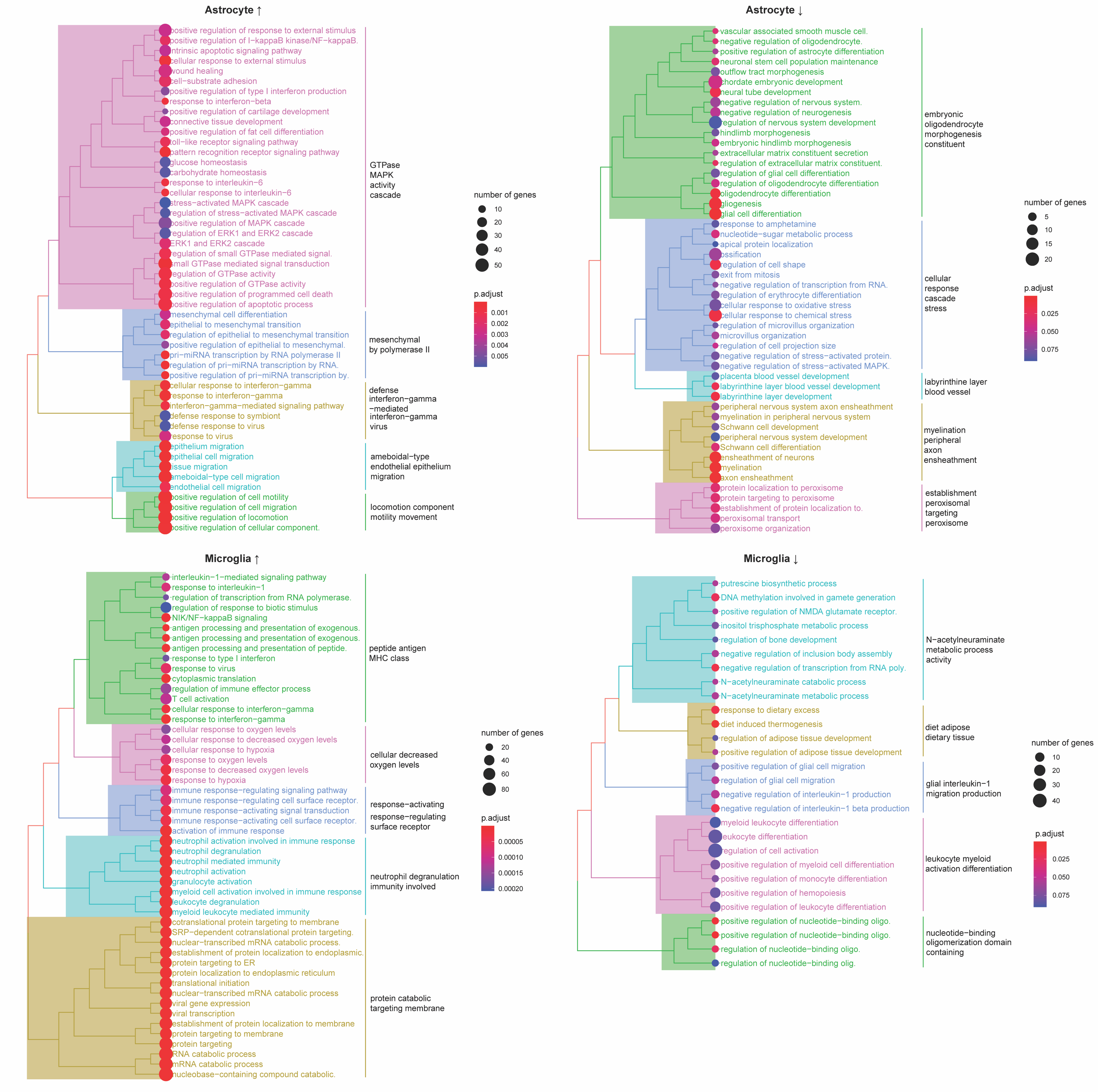

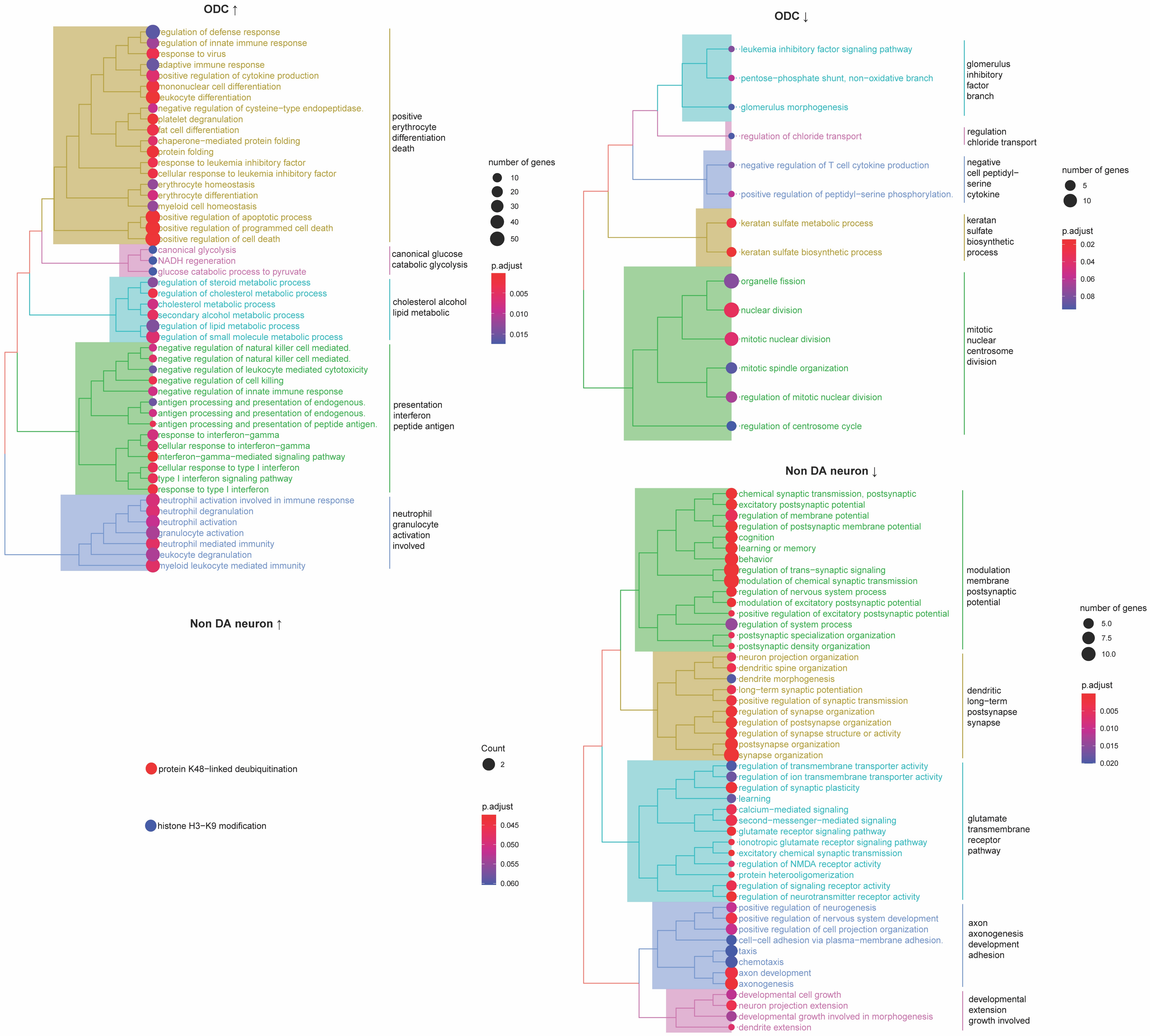

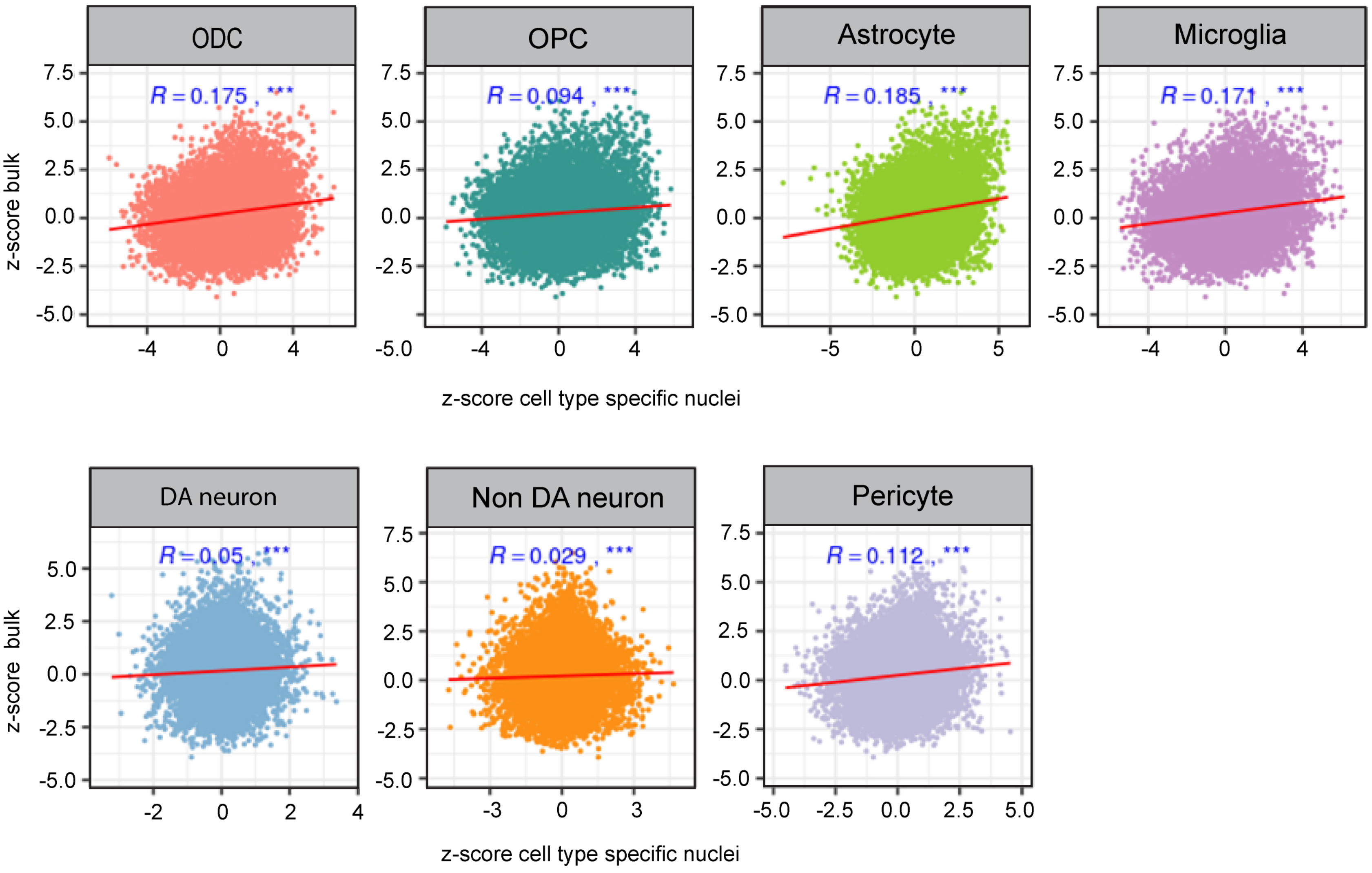

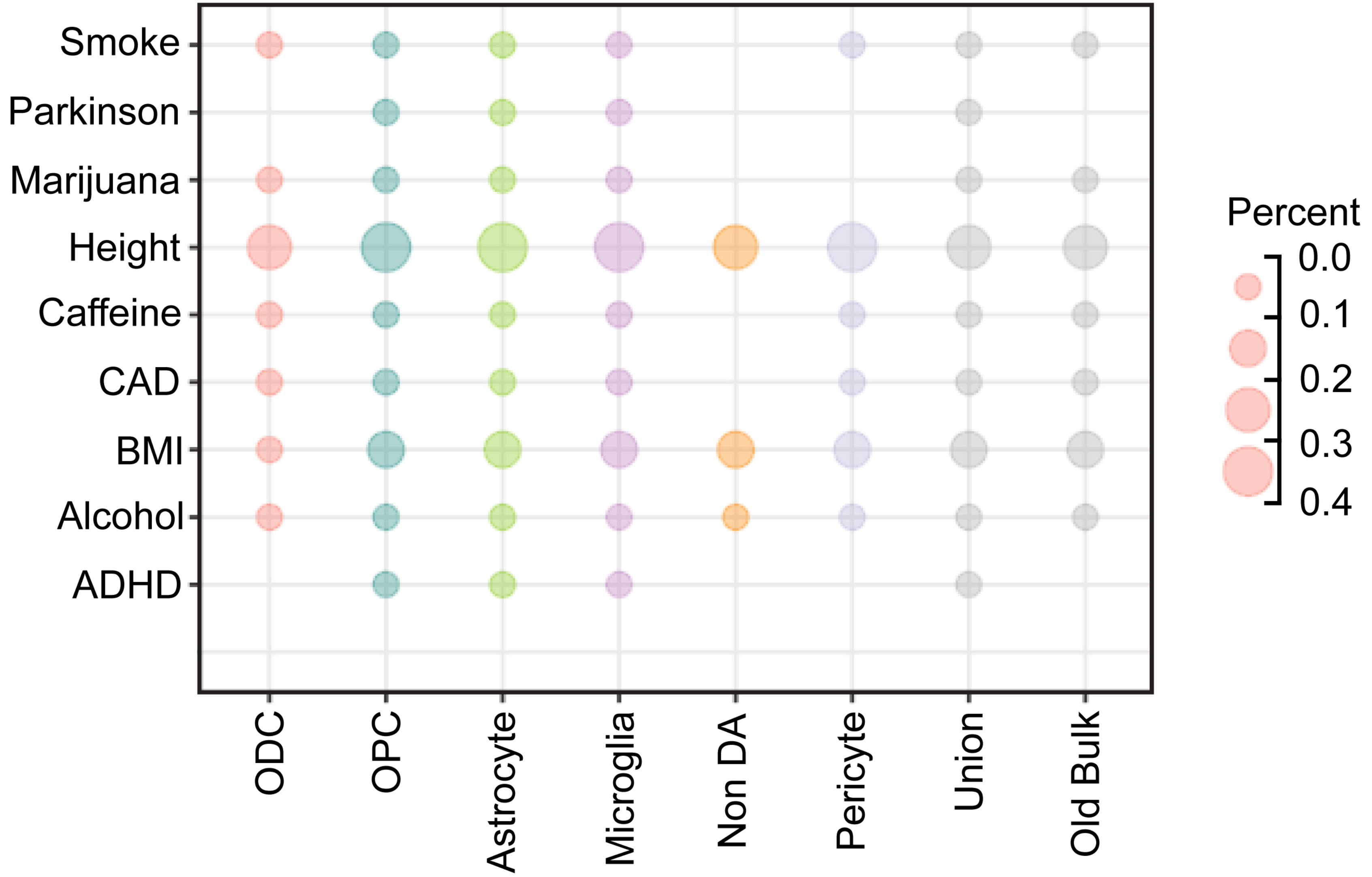

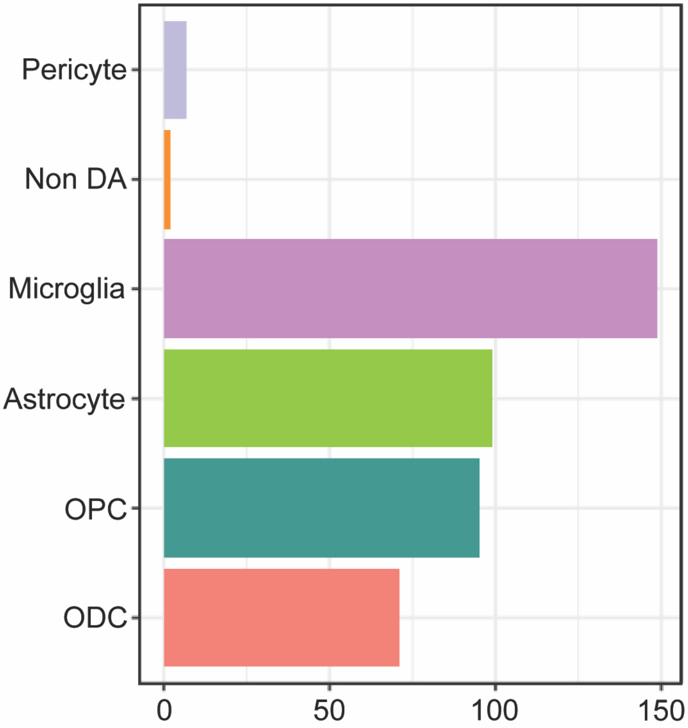
